# A Checking Model of Obsessive-Compulsive Disorder Based on the Hypothalamic Defensive System Against a Potential Threat

**DOI:** 10.1101/2022.11.01.514798

**Authors:** Noriko Horii-Hayashi, Kazuya Masuda, Taika Kato, Kenta Kobayashi, Mayumi Nishi

**Affiliations:** Department of Anatomy and Cell Biology, Faculty of Medicine, Nara Medical University, Kashihara, Nara 634-8521, Japan; Section of Viral Vector Development, National Institute for Physiological Sciences, Okazaki, 444-8585, Japan

**Author notes:** Corresponding author: Noriko Horii-Hayashi.

**Keywords:** Checking, Defensive behavior, Hypothalamus, Obsessive-compulsive disorder, Potential threat, Urocortin-3

## Abstract

**BACKGROUND:** Obsessive-compulsive disorder (OCD) is a psychiatric disorder characterized by obsessive thoughts and compulsive behavior. While some theories imply that OCD patients have cognitive biases and dysfunctional motivation regarding a potential threat, these views are not adequately supported by neurological evidence. Hypothalamic perifornical (PeF) urocortin-3 (UCN3) neurons are involved in defensive responses to a potential threat, and the activation of these neurons in mice induces repetitive and excessive checking and burying of novel objects. In this study, we evaluated the hypothesis that mice in which PeF UCN3 neurons are activated can serve as an OCD model.

**METHODS:** PeF UCN3 neurons were chemogenetically activated with clozapine-N oxide (CNO) in *Ucn3-Cre* mice. Marble-burying activity, repetitive/stereotypic behaviors in the homecage, and excessive responses to a novel object were measured as OCD-like behaviors. The effects of clinically used drugs for treating OCD on these behaviors were evaluated. The effect of CNO on neural activity in the cortico-striato-thalamo-cortical loop (which is regarded as an OCD circuit) was assessed with c-Fos immunolabeling.

**RESULTS:** CNO increased marble-burying activity, evoked homecage-specific repetitive/stereotypic behaviors that probably aimed to seal entrances, and induced repetitive and excessive checking and burying of novel objects. These behaviors were suppressed by selective serotonin-reuptake inhibitors but not by diazepam. CNO increased neural activity in the cortico-striato-thalamo-cortical loop.

**CONCLUSIONS:** These results indicated that mice whose PeF UCN3 neurons are activated can serve as a model of OCD, particularly as a checking model. This supports theories concerning the role of potential threats in the pathophysiology of OCD.

## INTRODUCTION

Obsessive-compulsive disorder (OCD) is a severe and disabling psychiatric disorder with a 2–3% lifetime prevalence (1). OCD is characterized by obsessive thoughts and compulsive behaviors that are expressed in a variety of ways, such as repetitive checking and excessive handwashing (2). The common understanding of OCD is that compulsive behavior caused by worrying obsessions is an attempt to relieve anxiety (2). However, the classification of OCD in the Diagnostic and Statistical Manual of Mental Disorders (Fifth Edition) has moved away from “anxiety disorders” into a new category of “obsessive-compulsive and related disorders” (3). This reclassification suggests that relationships between OCD and other anxiety disorders remain unclear. Numerous neuroimaging studies in people with OCD have indicated the hyperactivity of the cortico-striato-thalamo-cortical (CSTC) loop involving the orbitofrontal cortex (OFC) (4, 5), anterior cingulate cortex (ACC) (6–8), and caudate nuclei/striatum (Str) (4, 5, 9). In addition to the CSTC loop, the amygdala has been implicated in the pathophysiology of OCD (10–16).

The most effective treatment currently used for OCD is cognitive behavior therapy and high doses of selective serotonin-reuptake inhibitors (SSRIs) (17–19). Other non-serotonergic anxiolytics (e.g., benzodiazepines [BDZs]) are not effective in treating OCD (2), thus suggesting that SSRI efficacy is not dependent upon anxiolytic actions. Cognitive dysfunction is also involved in the pathophysiology of OCD (2, 20). Overestimation of threat and intolerance of uncertainty are widely accepted as key cognitive constructs underlying the maintenance of OCD symptoms (21–23). Such cognitive biases are particularly evident in patients with checking compulsions (24). OCD checkers are considered to have a lower threshold for perceiving ambiguous or uncertain situations as threatening (21).

Animals that encounter threatening situations exhibit defensive behaviors (25, 26). According to the Research Domain Criteria of the National Institute of Mental Health, threats can be categorized as either acute or potential. An acute threat is imminent and is an obvious risk to safety. Animals perceiving an acute threat (e.g., predators, intruders) exhibit defensive responses including fight, flight, and freezing. In contrast, a potential threat has ambiguous and uncertain risks. Animals encountering a potential threat, such as a novel object, exhibit risk assessment behavior (e.g., checking) (26–28). While a number of studies have investigated the neural mechanisms underlying defensive responses to acute threats (29–33), the mechanisms of defensive behaviors against potential threats remain unclear. Nevertheless, we recently reported that hypothalamic neurons in mice are involved in the modulation of defensive responses to potential threats (27).

Hypothalamic perifornical (PeF) urocortin-3 (UCN3) neurons are located between the paraventricular nucleus and fornix(27). These neurons largely project to the LS (LS) which is known to associate with moods, anxiety, and, psychiatric disorders (34–36). PeF UCN3 neurons respond to a novel object, and their activity is associated with the checking of novel objects. Chemogenetic activation of these neurons by Gq-based designer receptor exclusively activated by designer drugs (DREADD), evokes excessive checking of a novel object without affecting the anxiety level (27). Such activation also increased burying activity in the marble-burying test (27) which is regarded as a rodent repetitive/compulsive-like behavior (37–39).

As OCD patients have the cognitive biases of threat overestimation and uncertainty intolerance, we hypothesized that PeF UCN3 neurons are involved in the pathophysiology of OCD and that mice in which PeF UCN3 neurons are activated can serve as a model of OCD. To verify this hypothesis, we assessed such mice in terms of the following validities that are widely used for evaluating animal models for human psychiatric disorders: face (behavioral phenotype), predictive (therapeutic outcome), and construct (neural mechanism causing disorder-related phenotypes) (40–42). Regarding face validity, we have already reported that the activation of PeF UCN3 neurons induces repetitive and excessive checking and burying of novel objects and increases marble-burying activity (27). In the present study, we further investigate homecage-specific repetitive/stereotypic behaviors induced by the activation of PeF UCN3 neurons. To determine predictive validity, we examined the effects of SSRIs and BDZs on OCD-like behaviors caused by the activation of PeF UCN3 neurons. For construct validity, we investigated whether the activation of these neurons increases neural activity in the CSTC loop and limbic structures associated with negative valence including the amygdala and LS. Finally, we examined the effects of PeF UCN3 neuron activation on plasma corticosterone and adrenaline levels.

## METHODS AND MATERIALS

### Animals

All procedures for animal experiments were approved by the Animal Care Committee of Nara Medical University and were performed according to the National Institute of Health Guidelines and the Guidelines for Proper Conduct of Animal Experiments published by the Science Council of Japan. *Ucn3-Cre* mice were purchased from the Mutant Mouse Resource and Research Center (Stock #: 032078-UCD). Male mice aged 8–24 weeks were housed in a standard mouse cage with bedding material under standard laboratory conditions (23°C, 55% humidity, and a 12-h light-dark cycle: lights on at 8:00 a.m.) and had ad libitum access to food and water. For all experiments, mice were age-matched and randomly assigned to experimental groups to prevent a biased distribution of animals.

### Stereotaxic Surgery

Stereotaxic surgery was performed as described previously (27, 43). Adeno-associated virus (AAV) vectors [AAV(DJ): -hSyn-DIO-hM3D(Gq)-mCherry, 1 × 10^13^ copies/mL; AAV (DJ): -hSyn-DIO-mCherry, 1 × 10^13^ copies/mL) (Addgene, Watertown, MA) were injected into the PeF (250 nL/side, stereotaxic coordinate: Anterior-Posterior = −0.82 mm, Mid-Lateral = ± 0.47 mm, Dorsal-Ventral = 4.5 mm from the dura matter) at a flow rate of 100 nL/min. After surgery, mice were singly housed for 4 weeks and then used for the experiments. The accuracy of the injection site was checked by fluorescent microscopic observation in all mice that underwent surgery.

### Immunohistochemistry

Mice were anesthetized with sodium pentobarbital (100 mg/kg) and perfused transcardially with phosphate-buffered saline (PBS), followed by 4% paraformaldehyde in 0.1M phosphate buffer. Brains were post-fixed with the same fixative for 16 h at 4°C. Fifty-micrometer-thick sections cut using a vibratome (Microslicer; Dosaka, Kyoto, Japan) were permeabilized with PBS containing 0.3% Triton X-100 (PBST) and blocked with 5% normal horse serum in PBST. For immunofluorescent labeling, sections were incubated with primary antibodies diluted in the same blocking solution (guinea pig anti-c-Fos: 1:1000, Synaptic System, Goettingen, Germany; rabbit anti-UCN3: 1:200, Yanaihara, Shizuoka, Japan) for 2 days at 4°C. After three PBS washes, the sections were incubated with species-specific secondary antibodies conjugated to Alexa Fluor 488 ThermoFisher Scientific, Waltham, MA) for 2 h. After three additional PBS washes, the sections were mounted on glass slides and sealed with a mounting medium (ProLong Glass Antifade Mountant, Thermo Fisher Scientific). Fluorescent images were acquired and observed with a confocal microscope (FluoView 3000, Olympus, Tokyo, Japan).

Immunolabeling was performed with the diaminobenzidine (DAB)-avidin-biotin complex (ABC) method; sections were incubated with rabbit anti-c-Fos antibody diluted with the blocking solution (1:20000, #PC38, Calbiochem) for 2 days at 4°C. After three PBS washes, sections were reacted with biotinylated anti-rabbit IgG antibody (1:200, Vector Laboratory, Berlingem, CA) for 1 h. The sections were developed using a Vectastain ABC kit followed by a DAB Substrate Kit (Vector Laboratory), according to manufacturer instructions.

### c-Fos Expression Analysis

AAV-injected mice were intraperitoneally injected with saline or clozapine-N oxide (CNO) (2 mg/kg, Abcam, Cambridge, UK), returned to their homecage, and housed for 2 h. The mice were anesthetized with sodium pentobarbital (100 mg/kg) and fixed for immunolabeling of c-Fos as described above. Images were captured and observed with a microscope (BX-43, Olympus), and cell counting was performed with Flovel Image Filing System software (Flovel Corporation, Tokyo, Japan) by a trained experimenter blinded to experimental groups.

### Behavioral Testing

Behavioral testing was performed during the light phase from 9:30 to 13:00. Mice were transferred to a test room at least 20 min before commencing testing. All experiments were performed with a counterbalanced design across the sequence of treatments. Mice were subjected to a single test each day, with at least a 2-day interval between tests. Mice were intraperitoneally injected with saline or CNO (2 mg/kg). Either 20 mg/kg fluoxetine (FLX, Tokyo Chemical Industry Co., Ltd), 10 mg/kg escitalopram (ESC, Sigma-Aldrich), 1 mg/kg diazepam (DZP, Fujifilm Wako Pure Chemical Corporation), or a vehicle control was intraperitoneally injected 30 min before CNO injection.

The marble-burying test was performed 10 min after saline or CNO injection. Mice were allowed to move freely for 30 min in a standard laboratory cage containing 16 (4 × 4) glass marbles aligned on the surface of a bedding material of 4-cm thick. The number of buried marbles that were at least two-thirds covered by bedding material was counted.

The homecage test was performed after saline or CNO injection. The following homecage behaviors were measured over a 10-min period by a trained investigator blinded to experimental groups: pushing/piling bedding, rearing, and grooming. Locomotor activity was analyzed with TopScan LITE software (CleverSys Inc., Reston, VA).

The novel object test was performed after the homecage test. A novel object was placed along a wall of the homecage, and the behavioral response was serially recorded. The following items were used as a novel object, with a counterbalanced design across treatments: a ceramic house-shaped toy (3-cm height, 2-cm diameter), ceramic toothbrush stand (2.0 × 2. 0× 2.0 cm), bottle-shaped wood toy (4 × 5 × 1 cm), and a plastic ball (3-cm diameter). Mice were allowed to freely explore the object for 10 min. Sniffing was defined as having occurred when the distance between the nose and object was ≤ 1 cm. The following parameters were analyzed with TopScan LITE: number of sniffs, time spent sniffing, time spent staying on the object-side half of the cage, and locomotor activity. Burying was measured by a trained investigator blinded to experimental groups.

The new cage test was conducted 10 min after saline or CNO injection. Mice were allowed to freely move for 10 min in a standard mouse cage containing fresh bedding material. Rearing and digging were measured by a trained investigator blinded to experimental groups. Locomotor activity was analyzed with TopScan LITE.

### Corticosterone/Adrenaline Assay

Mice were injected with saline or CNO (2 mg/kg). Thirty minutes after injection, blood samples were collected from the facial vein with an animal lancet. Blood samples were centrifuged at 2,000 × *g* for 15 min at 4°C, and plasma was collected and stored at - 85°C. Corticosterone concentration was measured using an enzyme immunoassay kit (Yanaihara Institute, Fujinomiya, Japan), and adrenaline concentration was measured using an epinephrine enzyme-linked immunosorbent assay (ELISA) kit (Abnova corporation, Taipei, Taiwan).

### Statistical Analysis

GraphPad Prism 9 (GraphPad software incorporation, SanDiego, CA) was used to analyze data. The Mann–Whitney U test or Kruskal–Wallis test (followed by Dunnett’s post-hoc test) was used for non-paired comparisons. The Wilcoxon signed-rank test or Friedmann test (followed by Friedmann post-hoc multiple comparisons) was used for paired comparisons. *P*<0.05 indicated statistical significance.

## RESULTS

### Gq-DREADD Activation

Gq-DREADD AAV vector (-hSyn-DIO-hM3(Gq)-mCherry) or control vector (-hSyn-DIO-mCherry) was bilaterally injected into the PeF in *Ucn3-Cre* mice (Fig. 1A). Fluorescent microscopic observation showed that mCherry^+^ cells were distributed in the PeF between −0.58 and −0.94 mm from the bregma (Fig. 1B), which was consistent with our previous study findings (27). Immunofluorescent labeling of UCN3 showed overlap of mCherry^+^ cells and UCN3^+^ cells (Fig. 1C). To confirm the activation of mCherry^+^ cells using CNO, immunofluorescent labeling of c-Fos (a marker for activated neurons) was performed using saline- or CNO-injected mice. CNO (2 mg/kg) induced c-Fos expression in 96.0 ± 0.34% of mCherry+ cells (n = 5), whereas saline induced such expression in 2.12 ± 0.47% of mCherry+ cells (n = 5) (Fig. 1D) (Mann–Whitney U test: \**P*<0.05), thus indicating the activation of PeF UCN3 neurons by CNO.

**Fig. 1.**
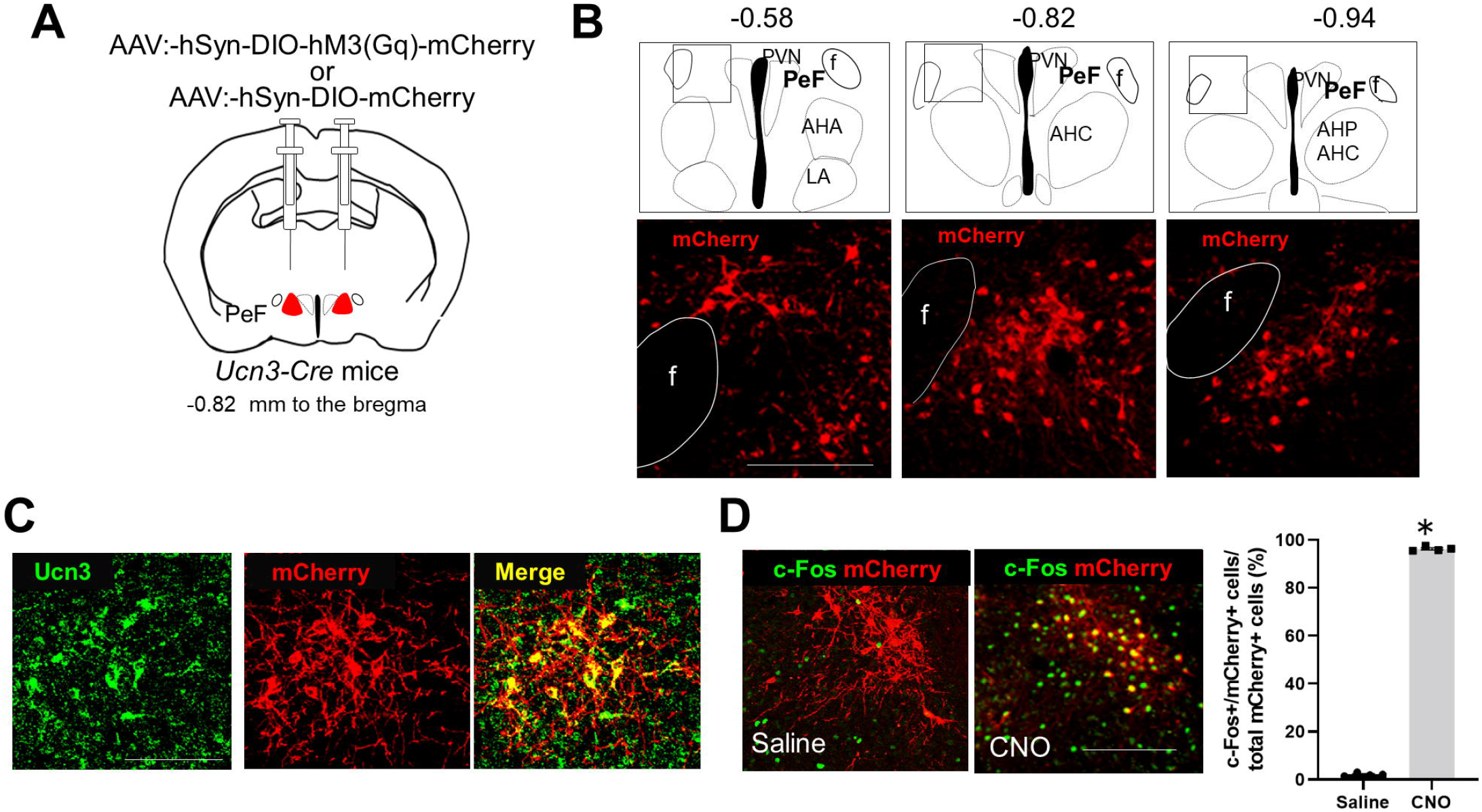
Chemogenetic activation of PeF UCN3 neurons. **(A)** Schematic diagram showing AAV injection into the PeF in *Ucn3-Cre* mice at −0.82 mm from the bregma. **(B)** Distribution of mCherry^+^ cells (red) in serial sections at −0.58 (left), −0.82 (middle), and −0.94 (right) mm from the bregma. Each square on the brain maps matches the microscopic field of fluorescent images below. **(C)** Immunofluorescent labeling of UCN3 in the PeF showing the overlap of UCN3^+^ cells (green) and mCherry^+^ cells (red). **(D)** Immunofluorescent labeling of c-Fos (green) in saline/CNO-injected mice. Graph shows the percentage of c-Fos^+^/mCherry^+^ cells to total mCherry^+^ cells (mean ± standard error: saline [2.1 ± 0.47 %], CNO [96.0 ± 0.34 %], n = 4, 4; Mann–Whitney test: U = 0, \**P*<0.05). Scale bars = 250 μm. AAV, adeno-associated virus; CNO, clozapine-N oxide; PeF, perifornical; UCN3, urocortin-3

### SSRI/DZP Effects on Marble-Burying Activity

CNO injection significantly increased marble-burying activity when compared with saline (saline: 10.1 ± 1.6 marbles, CNO: 14.6 ± 0.9 marbles, Wilcoxon signed-rank test: \**P*<0.05), whereas there was no significant difference between saline and CNO treatments in control AAV-subjected mice (saline: 9.57 ± 1.5 marbles, CNO: 8.9 ± 1.8 marbles, Wilcoxon signed-rank test, P = 0.59) (Fig. 2A). These results were consistent with that in our previous study (27). FLX (20 mg/kg) treatment 30 min before CNO injection in Gq-DREADD mice significantly reduced the increase in marble-burying activity caused by CNO, whereas DZP (1 mg/kg) did not show inhibitory effects (Fig. 2B) (n = 8, Friedmann test, analysis of variance [ANOVA]: *P*<0.01; Friedmann post-hoc test, vehicle/CNO(+) versus vehicle/CNO(−): *P*<0.05; vehicle/CNO(+) versus FLX/CNO(+): *P*<0.01; vehicle/CNO(+) versus DZP/CNO(+): *P*>0.99).

**Fig. 2.**
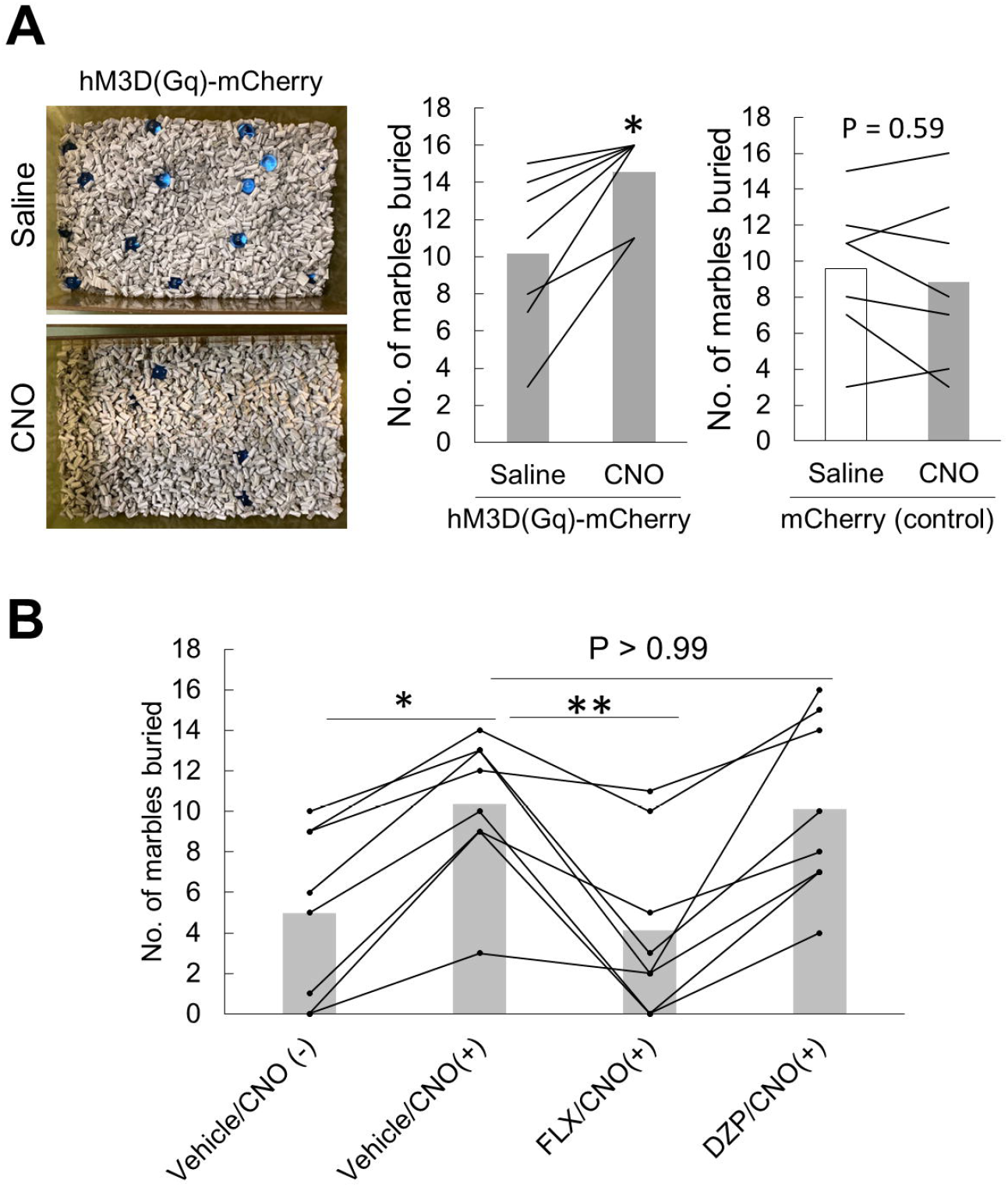
SSRI/DZP effects on marble-burying activity. **(A)** Pictures showing representative results of the marble-burying test using Gq-DREADD mice (top: saline; bottom: CNO). Graphs show the number of marbles buried after saline or CNO injection to Gq-DREADD mice (left) (n = 7, Wilcoxon signed-rank test, \**P*<0.05) and control AAV-subjected mice (right) (n = 7, Wilcoxon signed-rank test, P = 0.59). **(B)** Graph shows the numbers of marbles buried by Gq-DREADD mice injected with vehicle control/saline (Vehicle/CNO[−]), vehicle control/CNO (Vehicle/CNO[+]), FLX (20 mg/kg)/CNO (FLX/CNO[+]), and DZP (1 mg/kg)/CNO (DZP/CNO[+]). The order of drug administration was counterbalanced and randomly assigned to the mice (n = 8, Friedmann test, analysis of variance: *P*<0.01; post-hoc, \**P*<0.05). CNO, clozapine-N oxide; DREADD, designer receptor exclusively activated by designer drugs; DZP, diazepam; FLX, fluoxetine; SSRI, selective serotonin-reuptake inhibitor

### Homecage-Specific Repetitive/Stereotypic Behavior

CNO-injected Gq-DREADD mice showed characteristic repetitive/stereotypic behaviors in the homecage. For example, the mice pushed the bedding toward a wall and made a pile at the corner of the cage (Fig. 3A and Supplementary Movie 1). In the cage with a lid, CNO-injected mice reared and plugged the openings of the lid with bedding material (Fig. 3B and Supplementary Movie 2). To examine whether these behaviors were distinct from nest building, we performed the same experiment in the presence of cotton pads that had been provided as a nest material 7 days before the test. On the day of testing, we confirmed the presence of a cotton-made nest in all mouse cages used (data not shown). CNO-injected mice did not make contact with the cotton-made nest and showed similar behaviors to those observed in the absence of cotton pads (Fig. 3C and Supplementary Movie 3). Time-course observation showed that the pushing/piling of the bedding was evoked at approximately 10 min after CNO injection and lasted for approximately 20 min (Fig. 3D). In contrast, neither saline injection in Gq-DREADD mice nor CNO injection in control AAV-subjected mice resulted in this behavior (Fig. 3D) (n = 6, Kruskal–Wallis test followed by Dunnett’s post-hoc test, ANOVA: \*\**P*< 0.01; post-hoc test [Gq-mCherry × CNO versus Gq-mCherry × saline]: at 10 min [\**P*<0.05], at 15 and 20 min [\*\**P*<0.01], at 25 and 30 min [\*\*\**P*<0.001]). In addition to pushing/piling behavior, CNO also significantly increased rearing and locomotor activity when compared with saline (n = 10, Wilcoxon signed-rank test: pushing/piling, rearing, activity, \*\**P*<0.01); there was no significant difference in grooming (Fig. 3E) (n = 8, Wilcoxon signed-rank test: *P* = 0.44).

**Fig. 3.**
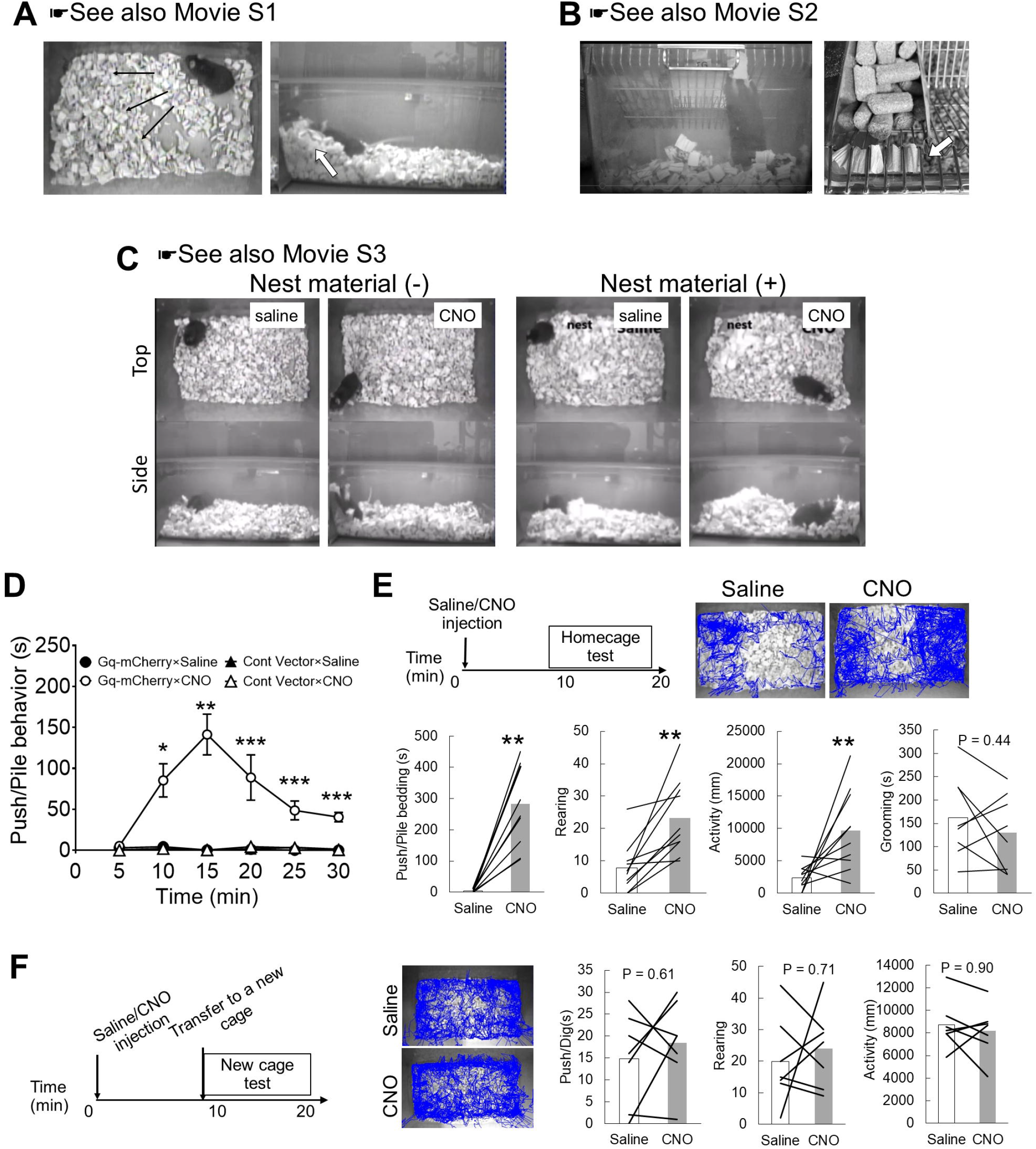
Homecage-specific repetitive/stereotypic behavior induced by CNO. **(A)** Appearance of CNO-injected Gq-DREADD mice in a homecage without a lid: the mouse pushed the bedding toward a corner/wall of the cage (black arrows) and made a pile (white arrow). (**B)** Appearance of a CNO-injected mouse plugging the openings of the cage lid with bedding. The white arrow indicates plugged bedding. **(C)** Top and side views of saline/CNO-injected mice in a homecage without (left) or with (right) cotton pads as a nest material. **(D)** Time-course changes in the behavior of pushing/piling bedding after saline or CNO injection (time 0) in mice injected with Gq-DREADD or control vector (n = 6, Kruskal–Wallis followed by Dunnett’s post-hoc test: \**P*<0.05, \*\**P*<0.01, \*\*\**P*<0.001 versus Gq-mCherry × saline). **(E)** A timeline of the homecage test among saline/CNO-injected mice. Representative images of nose-point tracking in saline/CNO-injected mice are shown. Graphs show the time engaged in pushing/piling bedding, the number of rearing events, locomotor activity, and the time engaged in grooming (n = 8–10, Wilcoxon signed-rank test: \*\**P*<0.01). **(F)** A timeline of the new cage test among saline/CNO-injected animals. Representative images of nose-point tracking in saline/CNO-injected animals are shown. Graphs show the time engaged in pushing/digging bedding, the number of rearing events, and locomotor activity (n = 7, Wilcoxon signed-rank test). CNO, clozapine-N oxide; DREADD, designer receptor exclusively activated by designer drugs

To examine whether the pushing/piling behavior was homecage-specific, we performed a similar experiment using the same standard cage containing fresh bedding (new cage) (Fig. 3F). As obvious piling behavior was not observed in the new cage, pushing or digging behavior was measured instead. However, there were no significant differences in pushing/digging, rearing, or locomotor activity (Fig. 3F) (n = 7, Wilcoxon signed-rank test: pushing/digging [*P* = 0.61], rearing [*P* = 0.71], locomotor activity [*P* = 0.90]). These results indicated that CNO induced pushing/piling behavior specifically in the homecage.

### SSRI/DZP Effects on Homecage Behavior

We next examined the effects of SSRIs and DZP on CNO-induced behavioral changes in homecages (Fig. 4A). Both FLX (20 mg/kg) and ESC (10 mg/kg) significantly suppressed CNO-induced pushing/piling behavior, without affecting rearing or locomotor activity, when compared with the vehicle control (Fig. 4B, 4C, and, Supplementary Movie 4) (FLX, n = 8, Wilcoxon signed-rank test: pushing/piling [*P*<0.01], rearing [*P* = 0.53], locomotor activity [*P* = 0.08]; ESC, n = 8, Wilcoxon signed-rank test: pushing/piling [*P*<0.01], rearing [*P* = 0.53], locomotor activity [*P* = 0.08]). In contrast, DZP had no significant effects on behaviors (Fig. 4D) (n = 7, Wilcoxon signed-rank test: push/piling bedding [*P* = 0.15], rearing [*P* = 0.93], locomotor activity [*P* = 0.58]).

**Fig. 4.**
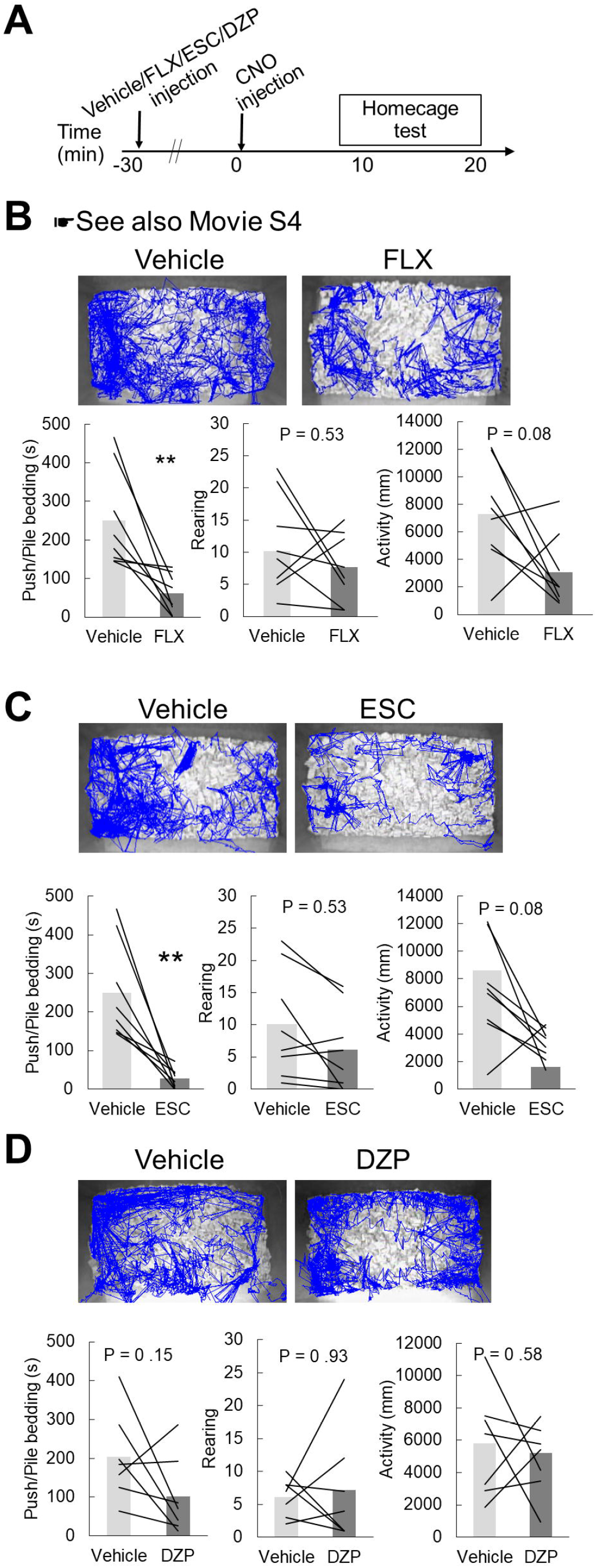
SSRI/DZP effects on CNO-induced homecage behaviors. **(A)** A timeline of the homecage test among mice treated with the vehicle control, FLX (20 mg/kg), ESC (10 mg/kg), or DZP (1 mg/kg) before CNO injection. **(B–D)** Representative images of nosepoint tracking in Gq-DREADD mice treated with FLX (B), ESC (C), or DZP (D) before CNO injection. Graphs show the time engaged in pushing/piling bedding, the number of rearing events, and locomotor activity (n = 8, Wilcoxon signed-rank test: \*\**P*<0.01). CNO, clozapine-N oxide; DREADD, designer receptor exclusively activated by designer drugs; DZP, diazepam; ESC, escitalopram; FLX, fluoxetine; SSRI, selective serotonin-reuptake inhibitor

### SSRI/DZP Effects on Responses to Novel Objects

After the homecage test, we next examined behavioral responses to a novel object in CNO-injected Gq-DREADD mice (Fig. 5A). The object sniffing zone was defined as a 1-cm region around the novel object, and the object-side zone was defined as half of the cage area on which the object was located (Fig. 5B). Consistent with that in our previous study (27), CNO injection significantly increased burying activity, duration of stay on the object side, the number of sniffs and time engaged in sniffing, and locomotor activity (Fig. 5C, 5D, and, Supplementary Movie 5) (n = 9, Wilcoxon signed-rank test: burying [\*\**P*<0.01], duration of stay on object-side zone [\*\**P*<0.01], number of sniffs [\*\**P*<0.01], time engaged in sniffing [\*\**P*<0.01], locomotor activity [\*\**P*<0.01]). Treatment with FLX or ESC 30 min before CNO injection (Fig. 5E) significantly reduced burying activity and duration of stay on the object side when compared with that with the vehicle control (Fig. 5F, 5G, and, Supplementary Movie 6) (n = 8, Wilcoxon signed-rank test: FLX, time engaged in burying [\*\**P*<0.01], duration of stay on object-side zone [\**P*<0.05], number of sniffs [*P* = 0.11], time engaged in sniffing [*P* = 0.078], locomotor activity [\*\**P*<0.01]; ESC: time engaged in burying [\*\**P*<0.01], duration of stay on object-side zone [\**P*<−0.05], number of sniffs [*P* = 0.11], time engaged in sniffing [*P* = 0.078], locomotor activity [\**P*<0.05]). DZP had no effects on behaviors (Fig. 5H) (n = 7, Wilcoxon signed-rank test: time engaged in burying [*P* = 0.90], duration of stay on object-side zone [*P* = 0.32], number of sniffs [*P* = 0.12], time engaged in sniffing [*P* = 0.16], locomotor activity [*P* = 0.80]).

**Fig. 5.**
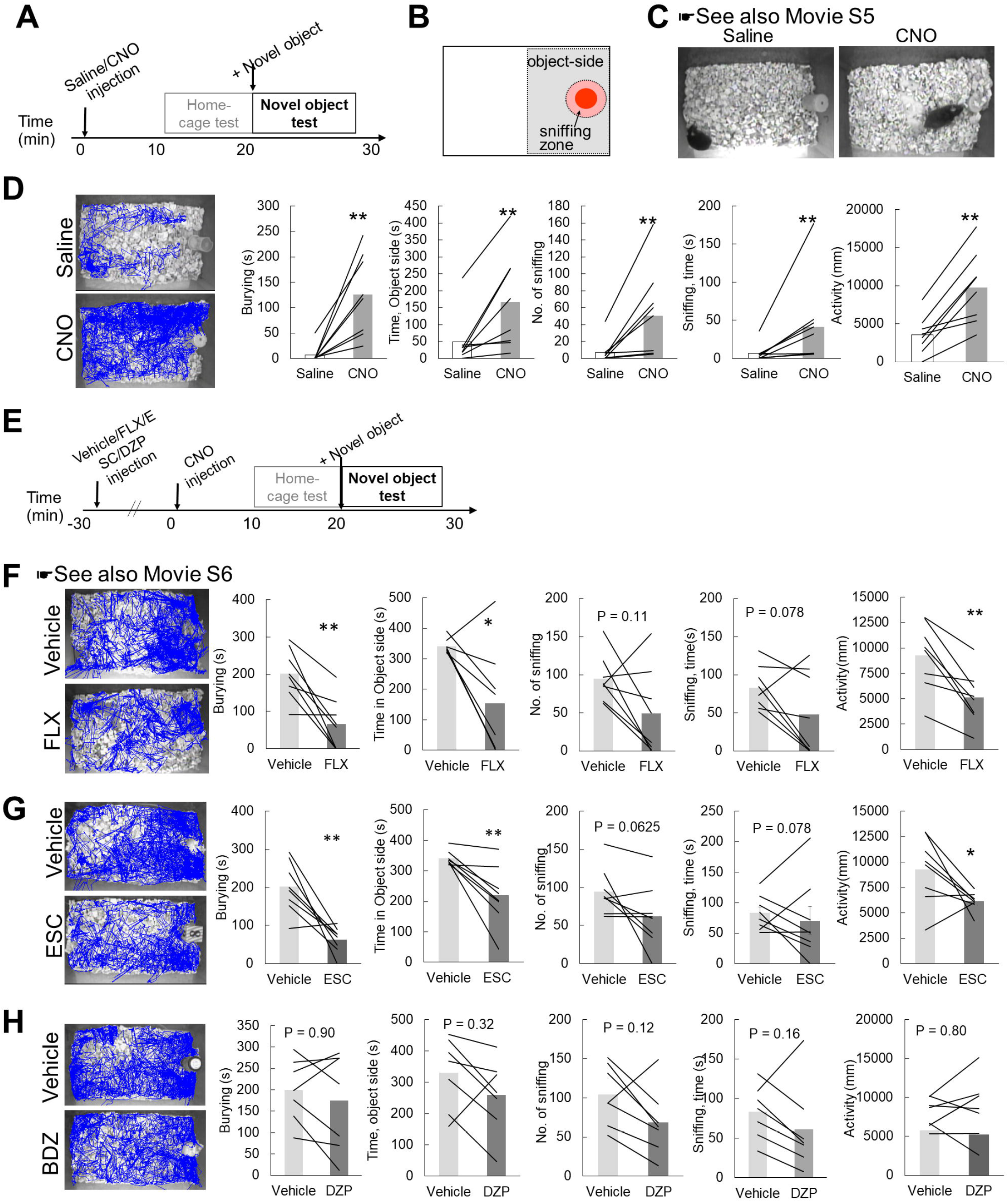
SSRI/DZP effects on CNO-induced responses to a novel object. **(A)** A timeline indicating measurement of behavioral responses to a novel object in a homecage using saline- or CNO-injected mice. A novel object was placed in a homecage after finishing the homecage test. **(B)** A 1-cm sniffing zone is shown around a novel object. The object-side area is defined as half of the cage area on which the novel object is located. **(C)** The appearance of saline- or CNO-injected Gq-DREADD mice. The latter exhibited vigorous burying of the novel object. **(D)** Representative images of nose-point tracking in saline- or CNO-injected mice. Graphs show the time engaged in burying, duration of stay on the object-side area, number of sniffs, time engaged in sniffing, and locomotor activity. **(E)** A timeline indicating measurement of behavioral responses to a novel object using mice treated with the vehicle control, FLX (20 mg/kg), ESC (10 mg/kg), and DZP (1 mg/kg) before CNO injection. (**F–H)** Representative images of nose-point tracking in FLX- (F), ESC- (G), and BDZ- (H) treated mice. Graphs show the time engaged in burying, duration of stay on the object-side area, number of sniffs, time engaged in sniffing, and locomotor activity (n = 7–8, Wilcoxon signed-rank test: \**P*<0.05 and \*\**P*<0.01). CNO, clozapine-N oxide; DREADD, designer receptor exclusively activated by designer drugs; DZP, diazepam; BDZ, benzodiazepine; ESC, escitalopram; FLX, fluoxetine; SSRI, selective serotonin-reuptake inhibitor

### Neural Activity in the CSTC Loop and Limbic Structures

We examined the effects of PeF UCN3 neuron activation on neural activity in the CSTC loop involving medial OFC (MO), ventral/lateral OFC LO/VO, ACC, and Str and, the limbic structures of the basolateral amygdaloid nucleus (BLA) and LS by c-Fos immunolabeling in saline/CNO-injected Gq-DREADD mice (Fig. 6A and 6B). All CNO-injected mice exhibited pushing/piling behavior in homecages (data not shown). A significant increase in the number of c-Fos^+^ cells was confirmed in the PeF (Fig. 6C) (n = 4, 5, Mann–Whitney U test: \**P*<0.05). CNO significantly increased the number of c-Fos^+^ cells in the MO, ACC, Str, BLA, and LS (Fig. 6D) (n = 4, 5, Mann–Whitney U test: PeF, MO, ACC, Str, LS, BLA [\**P*<0.05], LO/VO [*P* = 0.19]). These results indicated that the activation of PeF UCN3 neurons increased neural activities in the CSTC loop and the limbic structures associated with negative valence.

**Fig. 6.**
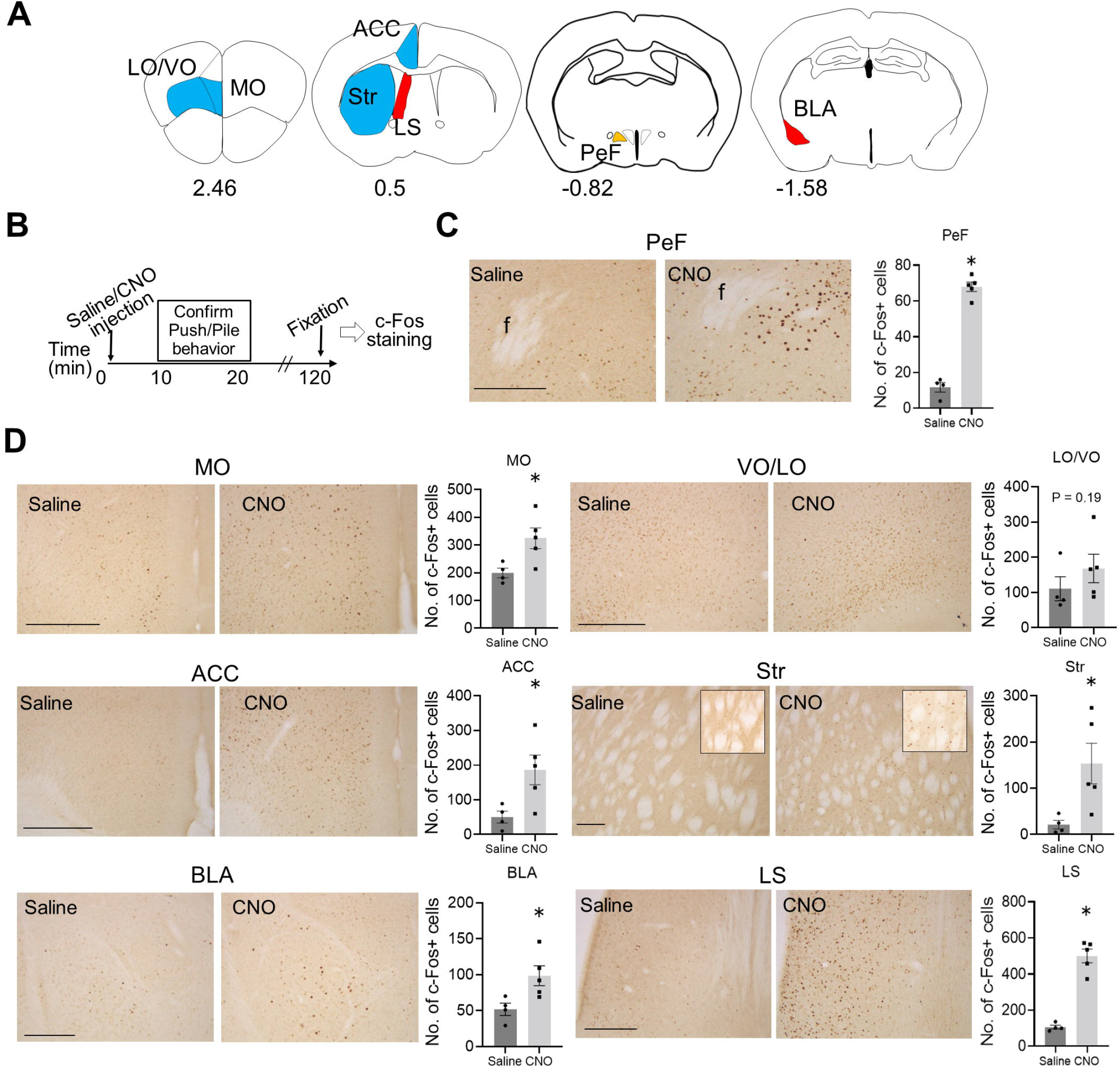
Neural activity in the CSTC loop and limbic structures. **(A)** Brain regions analyzed are indicated in blue (CSTC loop), red (limbic structures), and yellow (PeF). Numerical values indicate the distance from the bregma. **(B)** A timeline showing neural activity analysis via immunolabeling of c-Fos after evoking pushing/piling behavior with CNO in homecages. **(C and D)** Microscopic images showing c-Fos^+^ cells in the PeF (C), CSTC loop, and limbic structures (D) after saline/CNO injection. Graphs show the number of c-Fos^+^ cells (n = 4–5, Mann–Whitney test: \**P*<0.05). ACC, anterior cingulate cortex; BLA, basolateral amygdaloid nucleus; CNO, clozapine-N oxide; CSTC, cortico-striato-thalamo-cortical; LO/VO, lateral/ventral orbitofrontal cortex; LS, lateral septum; MO, medial orbitofrontal cortex; PeF, perifornical area. Scale bars = 500 μm.

### Corticosterone/Adrenaline

Blood samples were collected 30 min after saline/CNO injection, and plasma corticosterone and adrenaline concentrations were measured via an ELISA (Fig. 7A). All CNO-injected mice exhibited pushing/piling behavior in homecages (data not shown). There were no significant differences in either corticosterone or adrenaline concentrations between saline and CNO treatments (Fig. 7B) (n = 9, Wilcoxon signed rank test: corticosterone [*P* = 0.95], adrenaline [*P* = 0.82]).

**Fig. 7.**
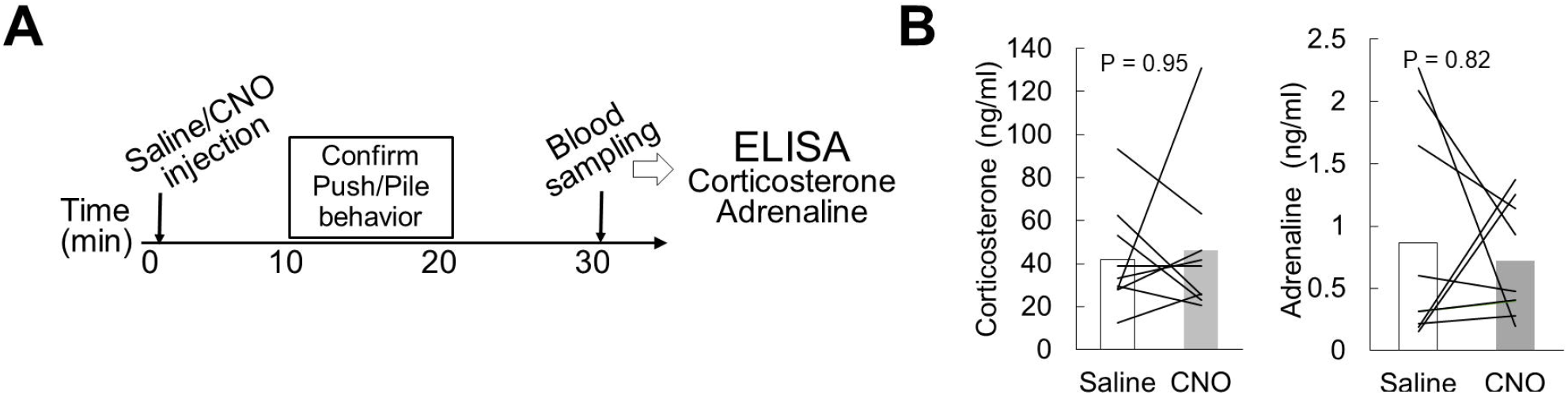
Plasma corticosterone and adrenaline concentrations. **(A)** A timeline showing the measurement of plasma corticosterone and adrenaline levels in blood samples collected after saline/CNO injection. **(B)** Graphs show plasma concentrations of corticosterone and adrenaline in saline/CNO-injected mice (n = 9, Wilcoxon signed rank test). CNO, clozapine-N oxide

## DISCUSSION

Animal models of OCD are useful as they provide insight into underlying genetic and neural mechanisms. They also facilitate the development of novel therapeutic agents and the elucidation of their mode of action (41, 44, 45). This study evaluated face, predictive, and construct validities using mice with activated PeF UCN3 neurons. Our results indicated that this mouse model fulfilled all three of these validities. In terms of face validity, PeF UCN3 neuron activation increased marble-burying activity, evoked homecage-specific repetitive/stereotypic behavior, and induced excessive checking of a potential threat (novel object). For predictive validity, we found that OCD-like behaviors caused by CNO were suppressed by SSRIs but not by DZP. In terms of construct validity, the activation of these neurons resulted in increased activity in the CSTC loop and the amygdala. Our results regarding plasma corticosterone and adrenaline levels exclude the possibility that the observed CNO-induced OCD-like phenotypes were caused through the activation of the hypothalamo-pituitary-adrenal axis and the sympathetic nervous system. Thus, we concluded that mice with activated PeF UCN3 neurons can serve as a useful animal model of OCD.

### Types of OCD model

Animal models of OCD are divided into multiple categories including genetic, pharmacological, and ethological models (41). The majority of current OCD genetic models have exhibited excessive self-grooming; these have included knockout mice for *HoxB8* (46), *Slitrk5* (47), *Sapap3* (48), and aromatase (49) as well as an optogenetic model stimulating the OFC-Str pathway (50). Among these models, the efficacy of SSRI (FLX) was confirmed in the optogenetic model and the *Slitrk5* and *Sapap3* models. Contrary to these models, the present model did not show excessive grooming in the homecage, nevertheless it showed increased activity in the CSTC loop. However, the CSTC loop is composed of multiple parallel and interconnected circuits that have individual functions including motor, cognitive, and emotional (44, 51). Thus, the present and previous studies suggest that compulsive-like grooming and checking are controlled by distinct or partially overlapping circuits within the CSTC loop. OCD-like compulsive checking is pharmacologically inducible by quinpirole, an agonist for the D2/D3 dopamine receptor (52, 53). While the efficacy of SSRIs has not been demonstrated in the quinpirole model, clomipramine has been shown to ameliorate quinpirole-induced compulsive checking (53). The model reported in the present study is the first genetic model to show a checking-like compulsion that fulfills all three validities.

### Homecage-specific Repetitive/Stereotypic Behavior

In rodent studies, there are two forms of defensive behavior related to home/territorial safety. The first is “aggression/fight,” which is a defensive response to imminent intruders. The second is “entrance sealing,” which was first described by Calhoun in a rat study (54). This is considered a defensive behavior against potential intruders. Calhoun observed that rats occasionally plugged entrances into their burrows with dirt, rocks, and vegetation from the inside of the burrow by repetitive movements of the nose/mouth and forepaws (54) (A video of entrance sealing can be viewed at https://youtu.be/xdgrD1VFx6k [4:16 to 4:54]). Notably, Calhoun categorized this behavior as a form of territorial defense and reported that the frequency of entrance sealing was increased by the state of lactation and a lower social rank (54). We speculate that the CNO-induced pushing/piling behavior in mice within homecages is analogous with entrance sealing in rats. In fact, in the presence of a cage lid, mice were observed to have plugged the openings of the lid with bedding material. Furthermore, there are three common features between entrance sealing and pushing/piling behavior: (i) territory/homecage-specific, (ii) occurrence within the cage/burrow, and (iii) repetitive movements of the nose/mouth and forepaws. The purpose of both behaviors is to keep the home safe by creating a barrier between the outside and inside of the cage/burrow that can repel intruders. This is similar to the human behavior of closing/locking the entrance of a house. Thus, CNO-induced repetitive pushing/piling in mice may be analogous to the repetitive checking of a door lock by people with OCD. Furthermore, the risk of OCD is higher in pregnant and postpartum women (55); this may be related to the observation of a higher frequency of entrance sealing among lactating female rats (54).

### Security Motivation System

The security motivation system is a theoretical model of OCD pathophysiology proposed by Szechtman and Woody (56, 57). This system comprises a set of biologically primitive behaviors (e.g., checking, washing) that are activated by a potential threat to self or intimate others (56, 57). While healthy people possess the security motivation system, its dysfunction leads to the failure to assess a potential threat and results in repetitive and excessive species-typical motor actions such as checking and washing (57). This theory is supported by our experimental evidence. The CSTC loop is involved in cognition, attentional control, motivation, motor control, and salience (58). Thus, PeF UCN3 neurons may drive a defensive motivation that assesses a potential threat via CSTC loop activation.

### Anxiety

The functional interrelationships among anxiety, obsessions, and compulsions are currently unclear. The inefficacy of BDZ-related anxiolytics in treating OCD casts doubt on the common explanation that compulsive behavior is an attempt to alleviate anxiety caused by worrying obsessions (2). In the present study, DZP failed to suppress CNO-induced OCD-like behaviors. Consistent with this result, our previous study showed that the activation of PeF UCN3 neurons had no effects on anxiety levels in the open-field, elevated-plus maze, or light–dark box tests (27). Thus, the present model may be optimal for dissecting the interrelationship between anxiety and compulsive behavior. While no prior clinical studies have implied that the hypothalamus is involved in the pathophysiology of OCD, the present findings suggest that hypothalamic neurons have a role in checking compulsion as well as dysfunctional responses to potential threats.

### Limitations

In the pharmacotherapy of OCD, a high dose of SSRIs should be administered for 8–12 weeks to attenuate symptoms (59). *Slitrk5*- and *Sapap3*-knockout OCD models, as well as the repeated stimulation of the OFC-striatal pathway (47, 48, 60), have consistently shown that chronic SSRI administration is necessary to suppress excessive grooming. However, the present model showed that acute SSRI administration was sufficient to suppress CNO-induced OCD-like behaviors. The exact therapeutic mechanisms of acute SSRI administration are currently unclear. Nevertheless, the effects of acute chemogenetic activation of PeF UCN3 neurons on the rest of the brain regions are considered more limited and transient when compared with that in the gene-knockout models and repeated stimulation model. Finally, the present study did not investigate the effects of PeF UCN3 neuron activation on cognitive functions such as reversal learning; this warrants investigation in future studies.

### Conclusions

The present study indicated that mice whose PeF UCN3 neurons are activated can serve as a model of OCD, particularly as a checking compulsion model. While there have been no clinical studies showing the involvement of the hypothalamus in the pathophysiology of OCD, the present findings support current theories concerning cognitive biases and dysfunctional security motivation regarding potential threats in the pathophysiology of OCD. The hypothalamus and UCN3 could be new targets for treating OCD.

## ACKNOWLEDGEMENTS

This study was supported by JSPS KAKENHI (Grant No. 22K03214 [to NH] and 19H03539 [to MN]) from the Japan Society for the Promotion of Science.

We thank Michiko Kitsuki and Mamiko Ozaki for their technical assistance.

## ARTICLE INFORMATION

From Nara Medical University (all authors).

NH conceived, designed, and performed the experiments and wrote the manuscript. KM and TK contributed equally to this work and analyzed behavioral data. MN supervised the work and contributed to the interpretation of data. All authors reviewed the manuscript draft and approved the final version of the manuscript to be published.

Address correspondence to Noriko Horii, Ph.D., at

